## Supplementary material for "Modular, robust and extendible multicellular circuit design in yeast": All SI figure and tables

### Supplementary Information

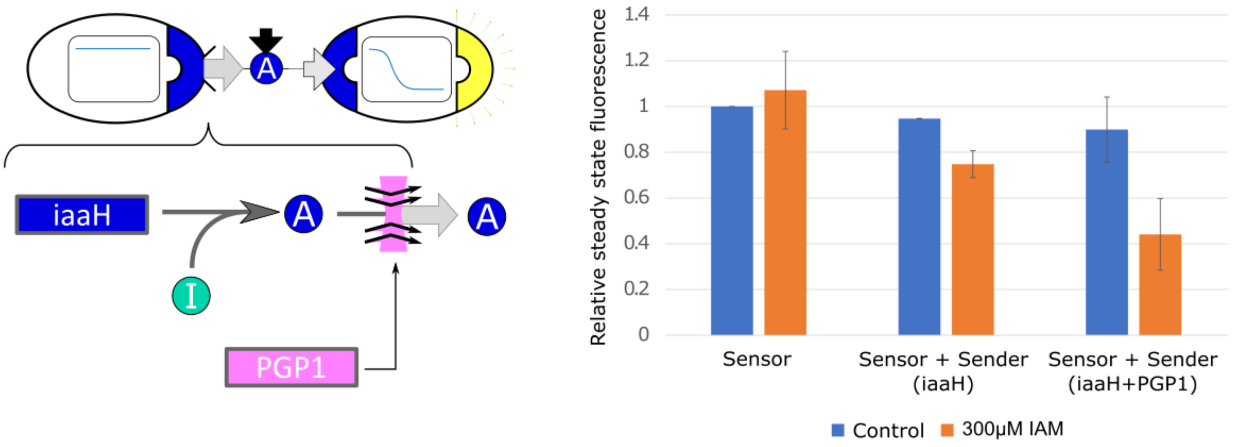


*Figure SI 1:* ***Effect of PGP1 on auxin secretion and sensing.*** *We integrated the auxin-pump PGP1 from A.thaliana into our yeast strains. Yeast strains expressing PGP1 have been shown to increase auxin efflux up to seven times [Geisler et al, 2005]. We observe a significant IAA-efflux increase in our sender/receiver co-culture, where the sender is constitutively synthesizing IAA from the IAM intermediate (I in the figure on the left). No improvements were detected with multiple integration of PGP1 (data not shown).*

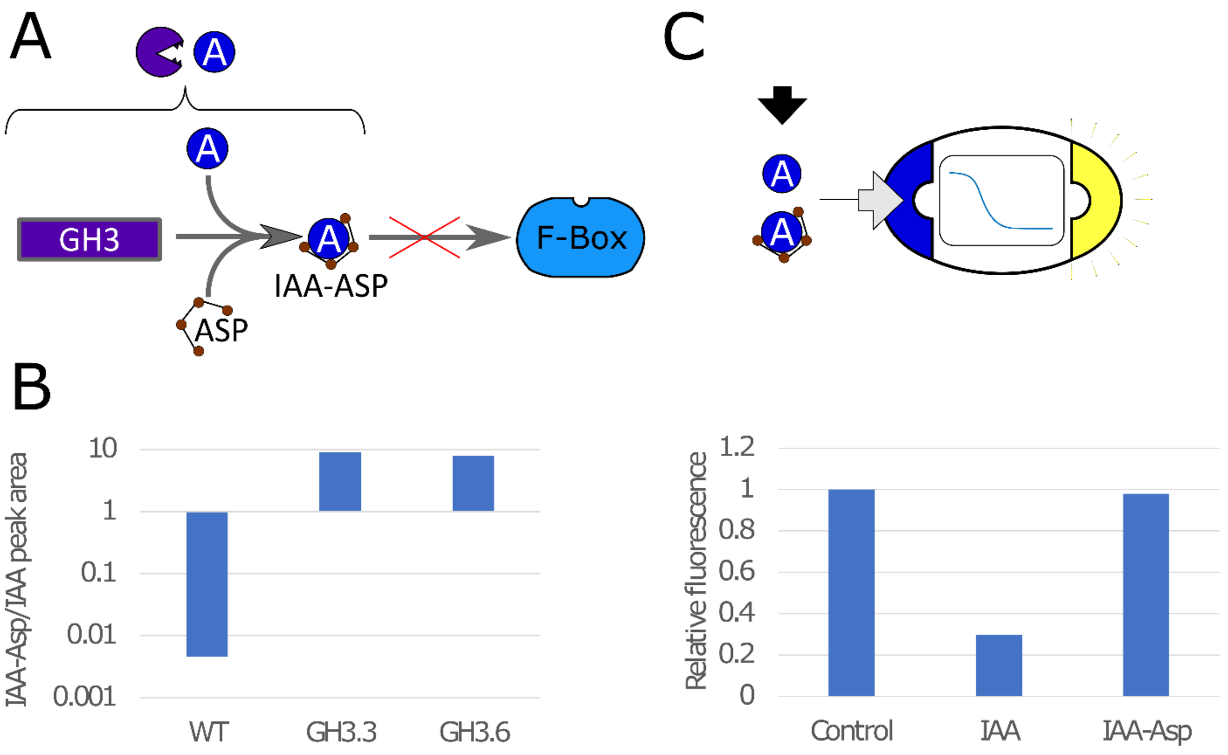


*Figure SI 2:* ***Auxin conjugation to Asp mediated by GH3 in yeast****. A) GH3 mediates binding of IAA to aspartic acid (Asp), making it not viable to bind to the F-box, which prevents auxin-induced degradation. B) Using mass-spectrometry (see Figure SI 3), we tested the effectiveness of GH3.3 and GH3.6 proteins of the GH3 family to bind IAA to Asp. Peak areas for IAA-Asp and IAA were detected with mass spectrometry and ratio is here reported for WT strain and strains expressing constitutively GH3.3 and GH3.6. The concentration of IAA-Asp is about 10 times higher than IAA for GH3-expressing strains. C) We tested for fluorescence response in strains that sense IAA and repress GFP for IAA-Asp, and we could not detect significant differences with respect to control. The bars are normalized so that Control is equal to 1.*

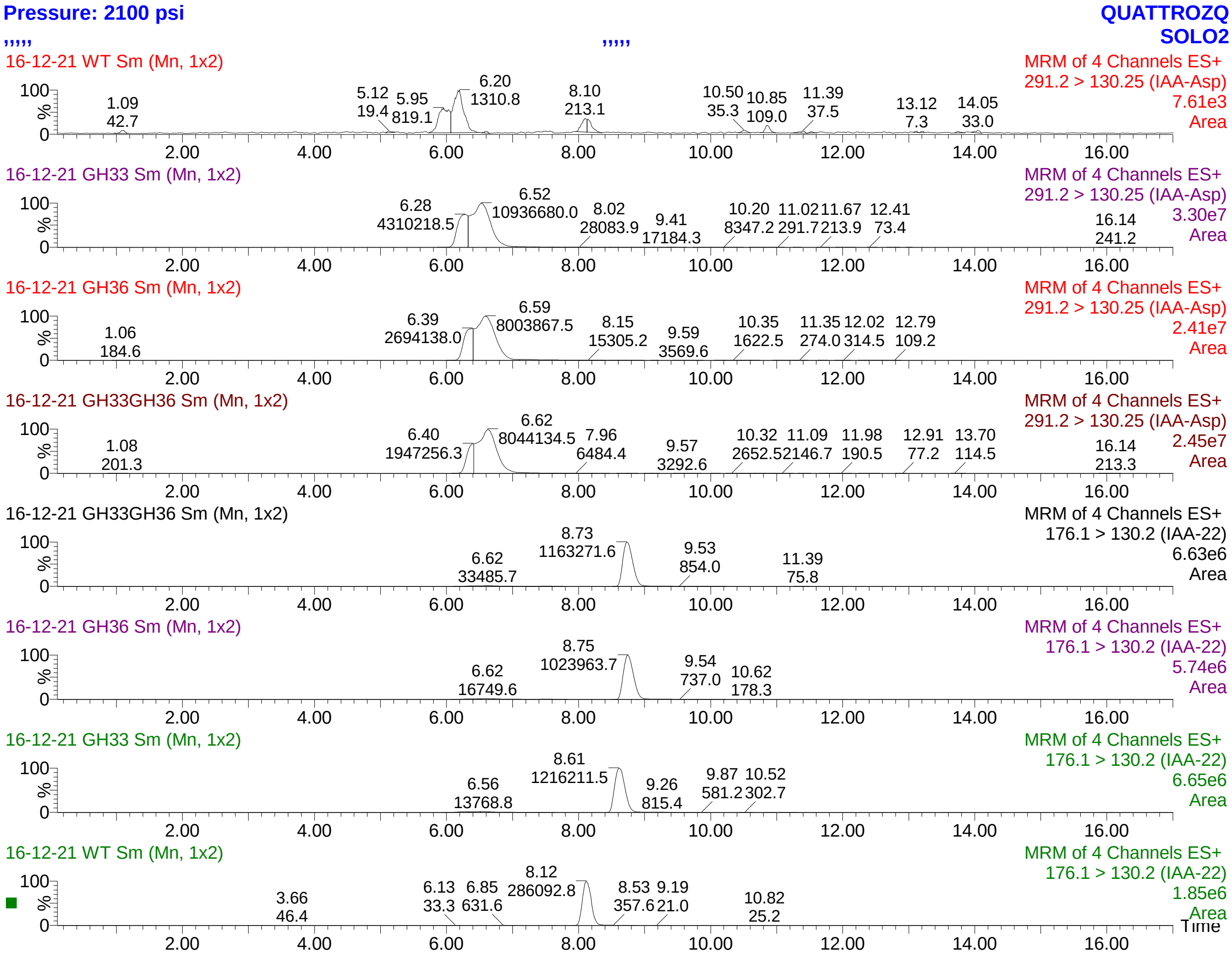


*Figure SI 3: **Mass spectrometry data for IAA-Asp synthesis from IAA in GH3-expressing strains.** We detected IAA and IAA-Asp peaks for WT strains and strains expressing GH3.3, GH3.6 and both proteins. Strains were exposed to 1µM IAA from low dilution till saturation (approximately 20 hours). Then cells were harvested, spun down and the supernatant was removed. After adding 250µL of methanol and boiled at 95°C for 15 minutes (vortexed twice during incubation) [\cit{lee2013expression }]. After spinning the cells down, we took the supernatant to perform IAA extraction through phase separation. We used the protocol outlined in \cit{kriechbaumer2016er} using ethyl acetate phase separation.*
*We expected a peak between 6-7 for IAA-Asp and between 8-9 for IAA. The label on the left-hand side shows the strain used, while the label on the right side shows the measured compound (IAA-22 or IAA-Asp). The number coming off the peak represent the area measurements used to generate plot B if Figure SI 2.*

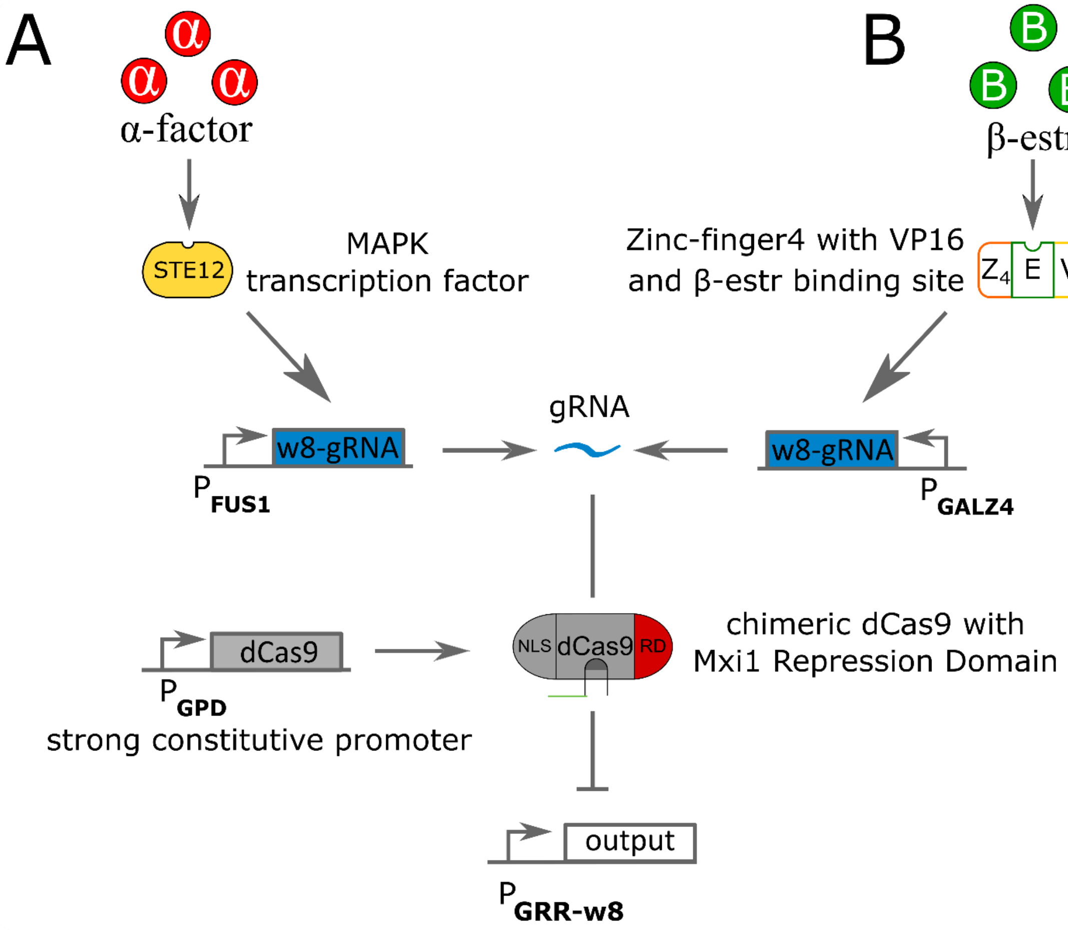


*Figure SI 4:* ***Circuit pathways for repressing strains using α-factor and β-estradiole as inputs. A)*** *The signaling molecule α-factor binds to the surface protein STE2, which activates a phosphorylation cascade that ends up activating the transcription factor STE12. Active STE12 then binds to the pFUS1 promoter, here rewired to synthesize gRNA (labelled w8 as in [Gander et al, 2018]). The gRNA binds to dCas9 fused with the Mxi1 repressing domain (RD) and a Nuclear Localization Signal. dCas9 is under a strongly constitutive promoter GPD. The gRNA-dCas9 complex binds to the pGRR-w8 promoter to stop transcription of the output gene.* ***B)*** *The signaling molecule β-estr penetrates the cell wall and binds to the chimeric protein Z4EV, constituted of a zinc-finger4 DNA-binding domain, a VP16 activation domain and binding pocket for β-estr. Upon β-estr binding, Z4EV induce transcription of the w8-gRNA gene under the pGALZ4 promoter. The rest of the pathway is identical to A)*

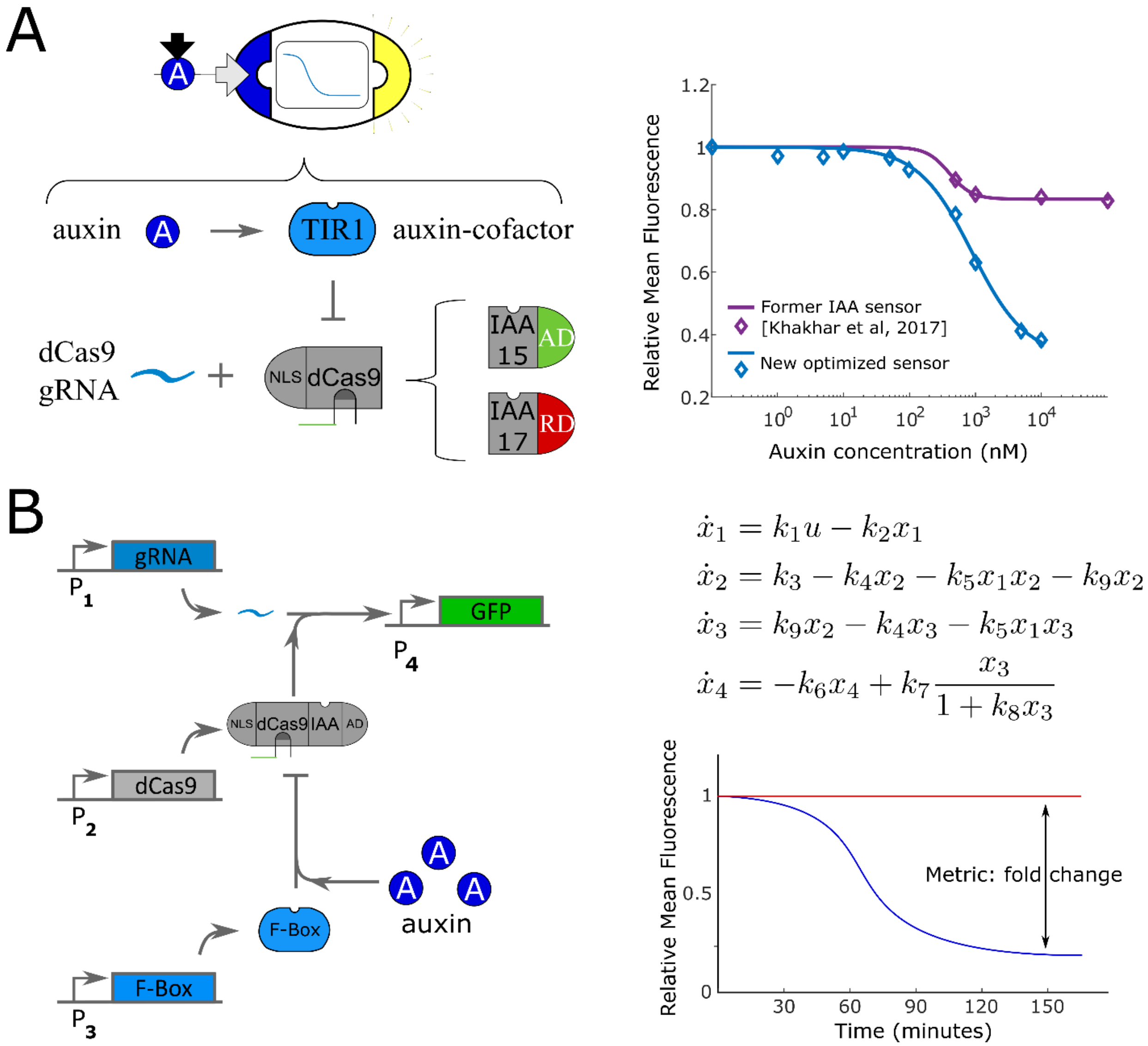


$$\dot{x}_1 = k_1 u - k_2 x_1$$
$$\dot{x}_2 = k_3 - k_4 x_2 - k_5 x_1 x_2 - k_9 x_2$$
$$\dot{x}_3 = k_9 x_2 - k_4 x_3 - k_5 x_1 x_3$$
$$\dot{x}_4 = -k_6 x_4 + k_7 \frac{x_3}{1 + k_8 x_3}$$

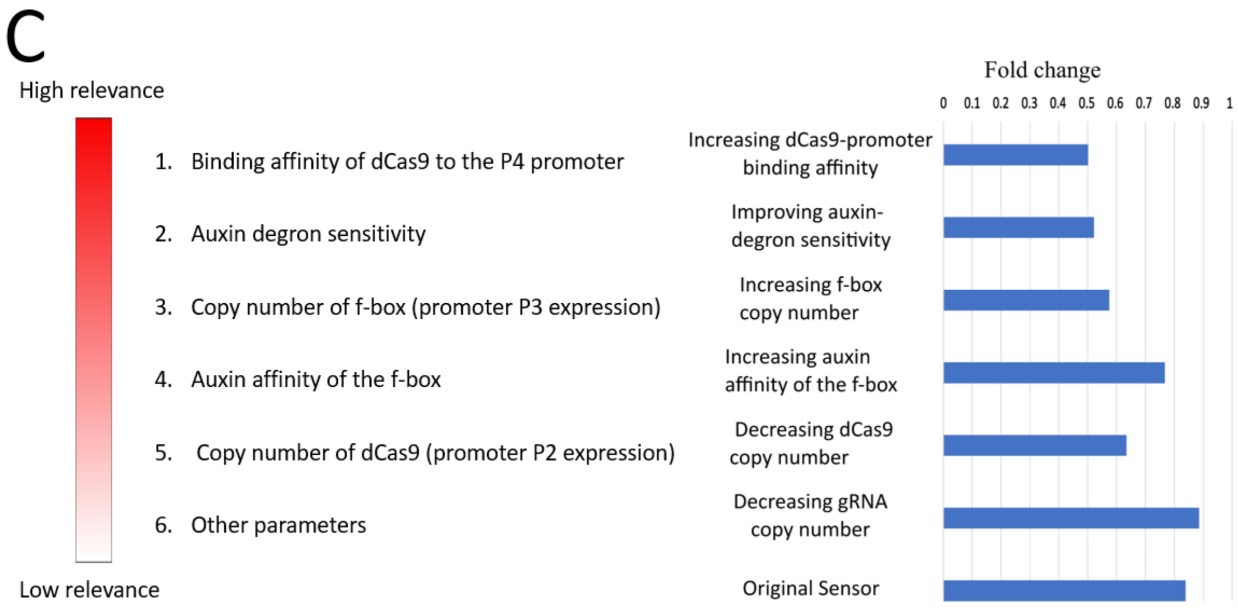


*Figure SI 5: **Pathway optimization for IAA-sensing strains**. A) IAA-sensing is mediated by degradation of a chimeric dCas9 with an activation or repression domain (AD or RD), which binds to a gRNA and to a promoter of interest in succession. When the promoter of interest drives GFP expression, we have a sensor strain. However, the repressing one proposed in \cit{Khakharetal} has low sensitivity and ON/OFF fold change (purple line in the steady-state plot). Hence, we optimized the pathway for higher sensitivity and fold change (blue line in the steady-state plot). Although here only the auxin-repressing pathway is depicted, a similar study was conducted for the IAA-activating pathway. B) To optimize the response, we minimized a metric defined as the L2-norm of the inverse of the fold-change. The fold change was computed by simulating the mechanistic model proposed in \cit{PierreJerome}, which is intuitive represented here, with constrained parameters $K_i$>0, i=1..9. C) We order the parameters according to their effect on the metric using sensitivity analysis, as shown here with next to the red bar. We then tested the effect of some of these perturbations in vivo and measured the fold-change between ON and OFF state (blue histogram to the right). The results agree with our predictions that increasing dCas9-promoter binding promoter and auxin-degron sensitivity would have the highest effect on fold-change. Driven by the model, we constructed a strain with three copies of TIR1_DM (f-box mutated for higher sensitivity), high affinity gRNA-promoter pair (w8 gRNA and pGRR-w8/pCYC1-w8 promoter, RD/AD) from \cit{Miles}, and IAA17/IAA15 (RD/AD) auxin degron for higher auxin sensitivity.*

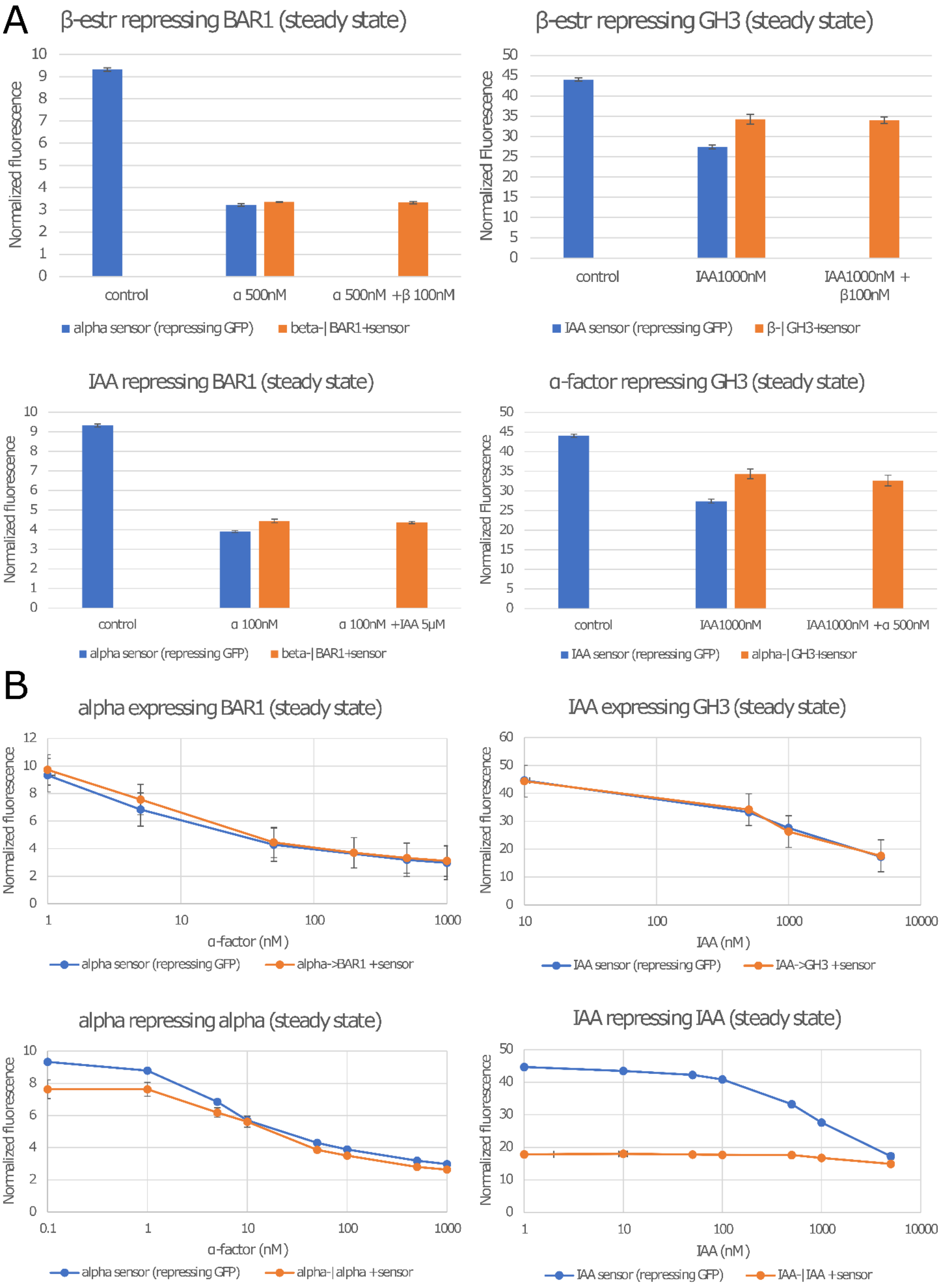
A
β-estr repressing BAR1 (steady state)
Normalized fluorescence
control
α 500nM
α 500nM +β 100nM
alpha sensor (repressing GFP)
beta-| BAR1+sensor
β-estr repressing GH3 (steady state)
Normalized Fluorescence
control
IAA1000nM
IAA1000nM + β100nM
IAA sensor (repressing GFP)
β-| GH3+sensor
IAA repressing BAR1 (steady state)
Normalized fluorescence
control
α 100nM
α 100nM +IAA 5μM
alpha sensor (repressing GFP)
beta-| BAR1+sensor
α-factor repressing GH3 (steady state)
Normalized Fluorescence
control
IAA1000nM
IAA1000nM +α 500nM
IAA sensor (repressing GFP)
alpha-| GH3+sensor
B
alpha expressing BAR1 (steady state)
Normalized fluorescence
α-factor (nM)
alpha sensor (repressing GFP)
alpha->BAR1 +sensor
IAA expressing GH3 (steady state)
Normalized fluorescence
IAA (nM)
IAA sensor (repressing GFP)
IAA->GH3 +sensor
alpha repressing alpha (steady state)
Normalized fluorescence
α-factor (nM)
alpha sensor (repressing GFP)
alpha-| alpha +sensor
IAA repressing IAA (steady state)
Normalized fluorescence
IAA (nM)
IAA sensor (repressing GFP)
IAA-| IAA +sensor

*Figure SI 6:* ***Strains constructed but not used in this study. A)*** *The four strains repressing expression of BAR1 or GH3 that are not positive feedbacks did not respond to their inputs significantly enough for us to include them in this study. They either do not affect the signaling molecule (the sensor response is identical to treatment whether the repressing strain is present of not), or they do not display a significant response range (no difference between the two orange bars).* ***B)*** *Top: these two negative feedback topologies do not significantly differ from the sensor alone in response to α-factor and IAA induction. Bottom: these strains present a significant functional range for negative feedback. Since, no circuits was built using these strains, they are not presented in the main manuscript.*

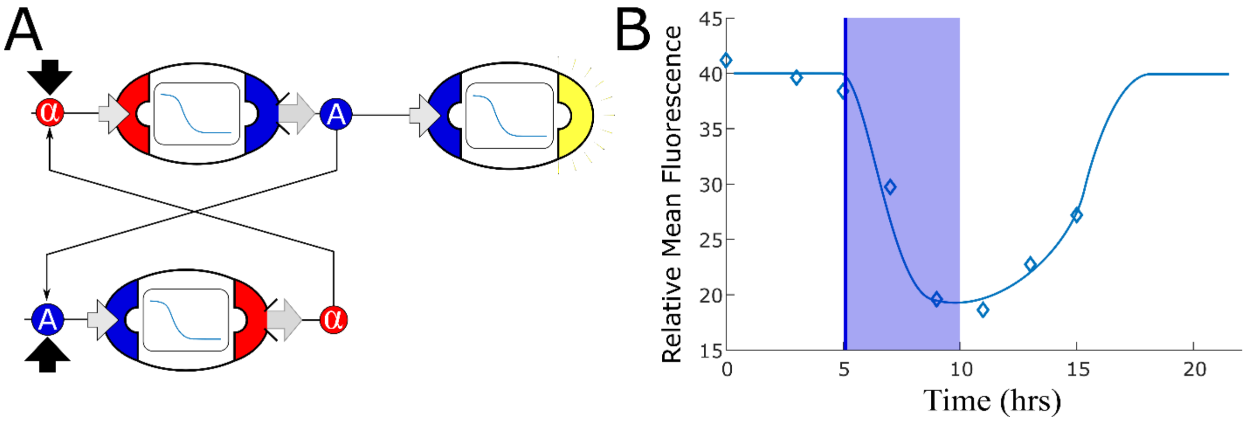


*Figure SI 7:* ***Mutually repressing strains do not generate bistability****. A) We tested a naïve realization of a potential bistable network using two strains that mutually repress each other's inputs: a strain that senses α-factor and represses auxin synthesis, and a strain that senses auxin and represses α-factor synthesis. B) We computed the optimal concentration of each of these strains using model simulation, but no combination generated two stable equilibria of α-factor and IAA concentrations. We experimentally tested the combination of these two strains with an auxin repression sensor strain to detect auxin concentration. Samples were taken every 2 hours along with dilution to restore the original cell concentration. After five hours, we inoculated 10μM IAA and diluted the samples of one third every 40 minutes, so that the IAA concentration became negligible after 5 hours (~150nM). The experimental data confirm that, while a lower value is reached when IAA is exogenously added, no stable equilibrium is reached and the system returns to the original equilibrium (low IAA concentration) after dilution. The diamonds are data, while the smooth line is the simulation; the transparent blue bar represents IAA inoculation starting at 5 hour, being reduced to below detection at hour 10.*

**Figure SI 8: Model derivation for the bistable switch**

Starting from the steady state expressions with general parameters:

$\alpha -| IAA$: $$x^{ss} = x_0 + \frac{a_0}{\varphi_0 + \alpha^{n_1}} \quad (1)$$

$\alpha \rightarrow GH3$: $$y^{ss} = y_0 + \frac{a_1 \alpha^{n_2}}{\varphi_1 + \alpha^{n_2}} \quad (2)$$

$IAA -| \alpha$: $$z^{ss} = z_0 + \frac{a_2}{\varphi_2 + IAA^{n_3}} \quad (3)$$

$IAA \rightarrow BAR1$: $$w^{ss} = w_0 + \frac{a_4 IAA^{n_4}}{\varphi_4 + IAA^{n_4}} \quad (4)$$

One obtains that the steady-state quantities of signaling molecules are:

$$IAA = \frac{IAA_0 + K_1 x^{ss}}{1 + K_3 y^{ss}} \quad (5)$$

$$\alpha = \frac{\alpha_0 + K_2 z^{ss}}{1 + K_4 w^{ss}} \quad (6)$$

where $IAA_0$ and $\alpha_0$ represents exogenous concentrations added to the mix, and $K_1$, $K_2$, $K_3$, and $K_4$ are the initial concentrations of each cell type (where K=1 is the standard value).

Substituting the above expressions (1)-(4) in (5) and (6) and re-labelling the parameters, one obtains:

$$IAA = \frac{a_1 IAA_0 + a_2 IAA_0 \alpha^{n_1} + a_3 IAA_0 \alpha^{n_2} + IAA_0 \alpha^{n_1+n_2} + K_1(c_1 + c_2 \alpha^{n_1} + c_3 \alpha^{n_2} + c_4 \alpha^{n_1+n_2})}{b_0 + b_1 K_3 + \alpha^{n_1}(c_1 + c_2 K_3) + \alpha^{n_2}(d_1 + d_2 K_3) + \alpha^{n_1+n_2}(1 + eK_3)} \quad (7)$$

$$\alpha = \frac{f_1 \alpha_0 + f_2 \alpha_0 IAA^{n_3} + f_3 \alpha_0 IAA^{n_4} + \alpha_0 IAA^{n_3+n_4} + K_2(h_1 + h_2 IAA^{n_3} + h_3 IAA^{n_4} + h_4 IAA^{n_3+n_4})}{g_0 + g_1 K_4 + IAA^{n_3}(j_1 + j_2 K_4) + IAA^{n_4}(l_1 + l_2 K_4) + IAA^{n_3+n_4}(1 + mK_4)} \quad (8)$$

If one simplifies the expression above considering only the highest and lowest terms, one obtains:

$$IAA = \frac{a_1 IAA_0 + IAA_0 \alpha^{n_1+n_2} + K_1(c_1 + c_4 \alpha^{n_1+n_2})}{b_0 + b_1 K_3 + \alpha^{n_1+n_2}(1 + eK_3)} \quad (9)$$

$$\alpha = \frac{f_1 \alpha_0 + \alpha_0 IAA^{n_3+n_4} + K_2(h_1 + h_4 IAA^{n_3+n_4})}{g_0 + g_1 K_4 + IAA^{n_3+n_4}(1 + mK_4)} \quad (10)$$

Finally, if one relabels the parameters using $\tilde{n}_1 = n_1 + n_2$ and $\tilde{n}_2 = n_3 + n_4$, obtains the same expression as in Figure 3C, Model 3.

Expressions (7) and (8) have then been fitted to simulation data of the correspondent strain combinations for different α-factor and IAA concentration to estimate the values of the parameters. Accordingly, the new exponents are: $\tilde{n}_1 = 2.003$ and $\tilde{n}_2 = 1.78$.

| Equation | $a_1$ | $\tilde{n}_1$ | $c_1$ | $c_4$ | $b_0$ | $b_1$ | $e$ |
|---|---|---|---|---|---|---|---|
| IAA | 101.90 | 2.003 | 4.79e+03 | 6.16e+06 | 494.41 | 3.21e-12 | 17.59 |

| Equation | $f_1$ | $\tilde{n}_2$ | $h_1$ | $h_4$ | $g_0$ | $g_1$ | $m$ |
|---|---|---|---|---|---|---|---|

| | | | | | | | |
|---|---|---|---|---|---|---|---|
| α | 428.28 | 1.78 | 1.65e+04 | 3.28 | 1.406e+03 | 3.59e+04 | 233.68 |

To study the existence of two stable solutions, we solved system (9) + (10) numerically with different values of the parameters $K_i$ and identified the area when multiple solutions arise.

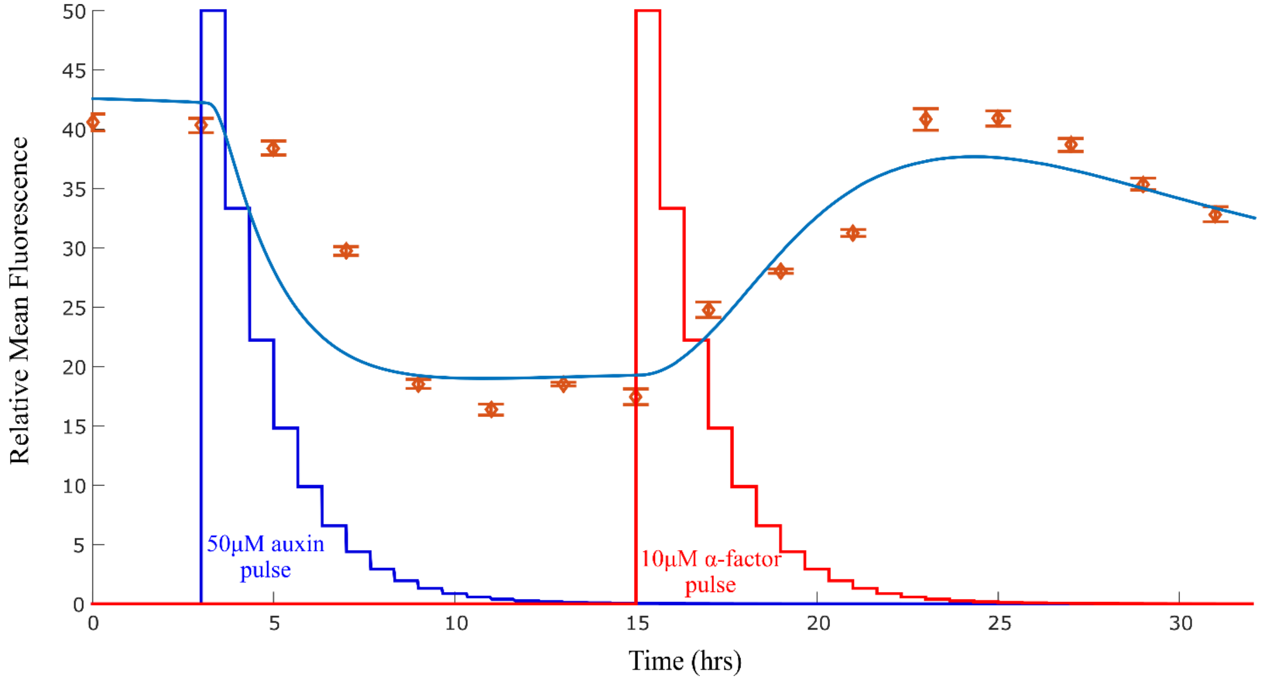


*Figure SI 9: **Model explanation for the stable-to-unstable shift**. The experimental data for the bitable switch constructed as in Figure 3C appear to exit its stable orbit after hour 25, as one can see from the steady decrease in fluorescence. One potential explanation is that the ratio between the strains is no longer maintained due to small difference in strain growth rate that we did not detect over 10 or 12 hours in this study. To test this hypothesis, we simulated the system and allowed the strain that synthesis BAR1 to slightly increase in concentration over time (reaching 0.8 at time 25 hour from a starting concentration of 0.7). With this minor correction, the simulation predicts loss of stability as observed in the data, with the three equilibria becoming a single stable equilibrium with high IAA and low α-factor concentration.*

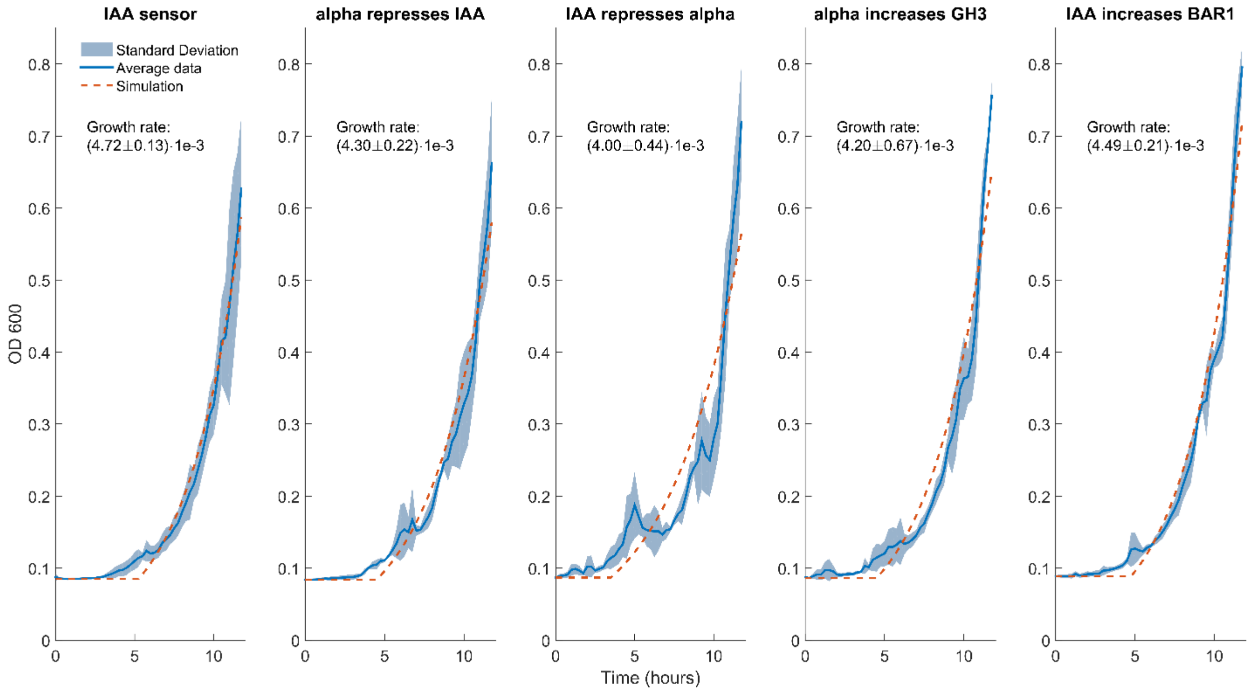


*Figure SI 10: Growth rate of the five strains in the bistable switch. Data collected with a plate reader over the course of 12 hours, 3 experimental repeats for each of the five strains in the bistable switch topology. Data were collected as OD600, sampling every 15 minutes at 30C, starting from an initial concentration of 30 events/µl. Growth rate was estimated fitting an exponential curve and the value here reported is average and standard deviation over fitting each of the three replicates. To account for the lower sensitivity threshold of the plate reader, each time series was set to be the minimum detectable till a 25% increase was detected.*

**Figure SI 11: Normalization procedure for logic gate vectors**

Each simulation ran 3 differential equations for each node, plus pooling parameters that represent the overall concentration of auxin and alpha-factor accounting for exogenous inputs, strain secretion, and BAR1 or GH3 synthesis, the two latter quantities being dependent on the strain selection. To automate the simulations, we represented each strain as a vector of parameters: the first 9 entries being the differential equation parameters, followed by one-hot encoding vector of size 3 to represent the sensed input ([beta-estr, alpha-factor, auxin]), and a one-hot encoding vector of size 4 to represent the secreted output ([alpha-factor, auxin, BAR1, GH3]). The last entry of the vector is a 0 for repressing strains and 1 for activating strains. Along the selected strains, we simulated the 8 sensor strains (Figure 1C, D, E) as system outputs.

We then grouped the simulations according to the input entries of the logic tables obtained from the 3 possible permutations of the inputs ([beta,alpha],[beta,auxin],[auxin,alpha]). These groups are shaped as a vector with 4 entries that correspond to the input states ([0 0], [1 0], [0 1], [1 1]). Within each group, we normalized each vector by their minimum value and subtracted 1 (the minimum becoming 0). This normalization scheme allowed comparison between different topologies accounting for differences in the sensor strains. At the end of this process, we had 120 vectors resulting from the 2-node networks, 560 from the 3-node networks, and 1820 from the 4-node networks.

### AND gates

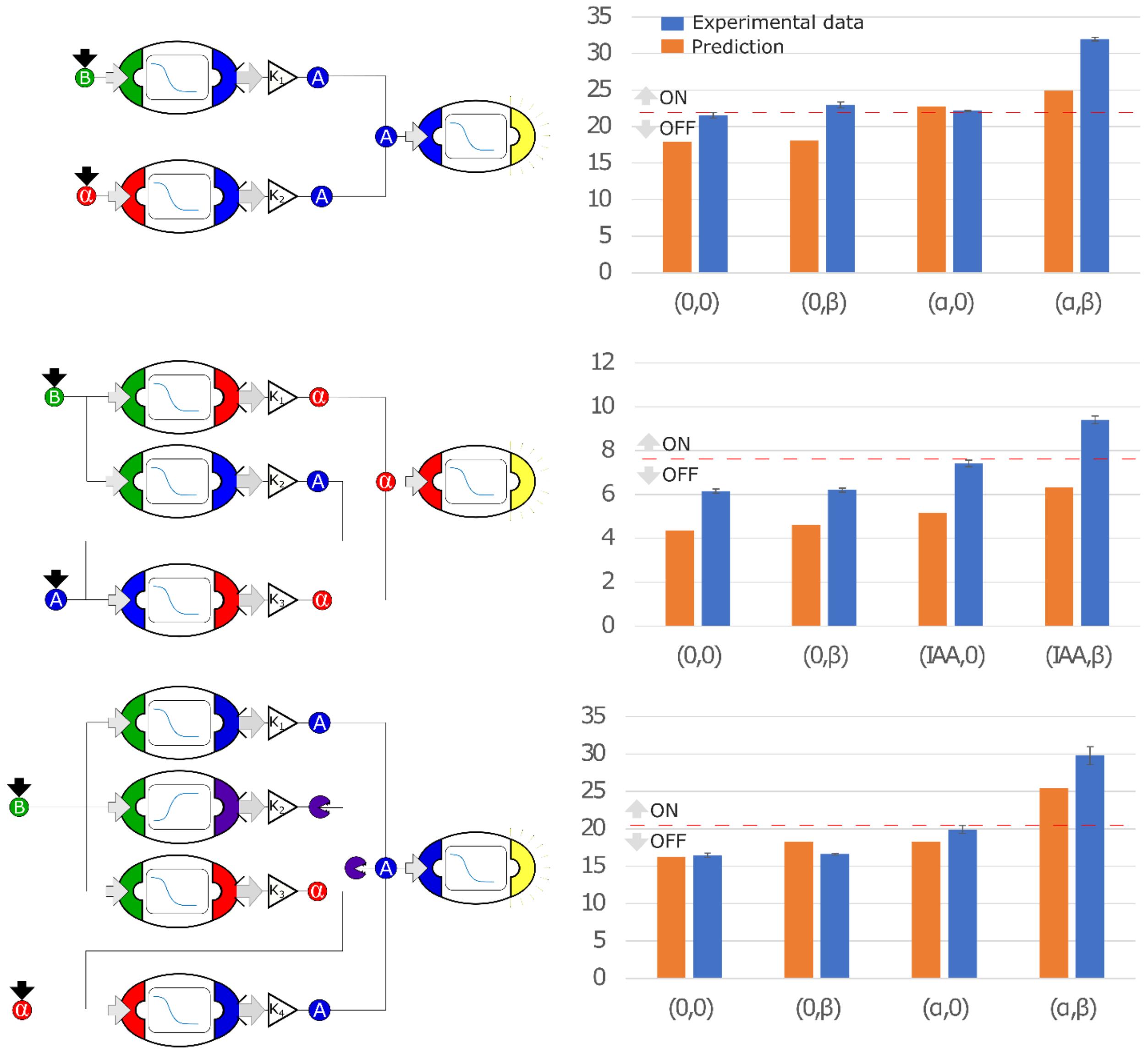

### OR gates

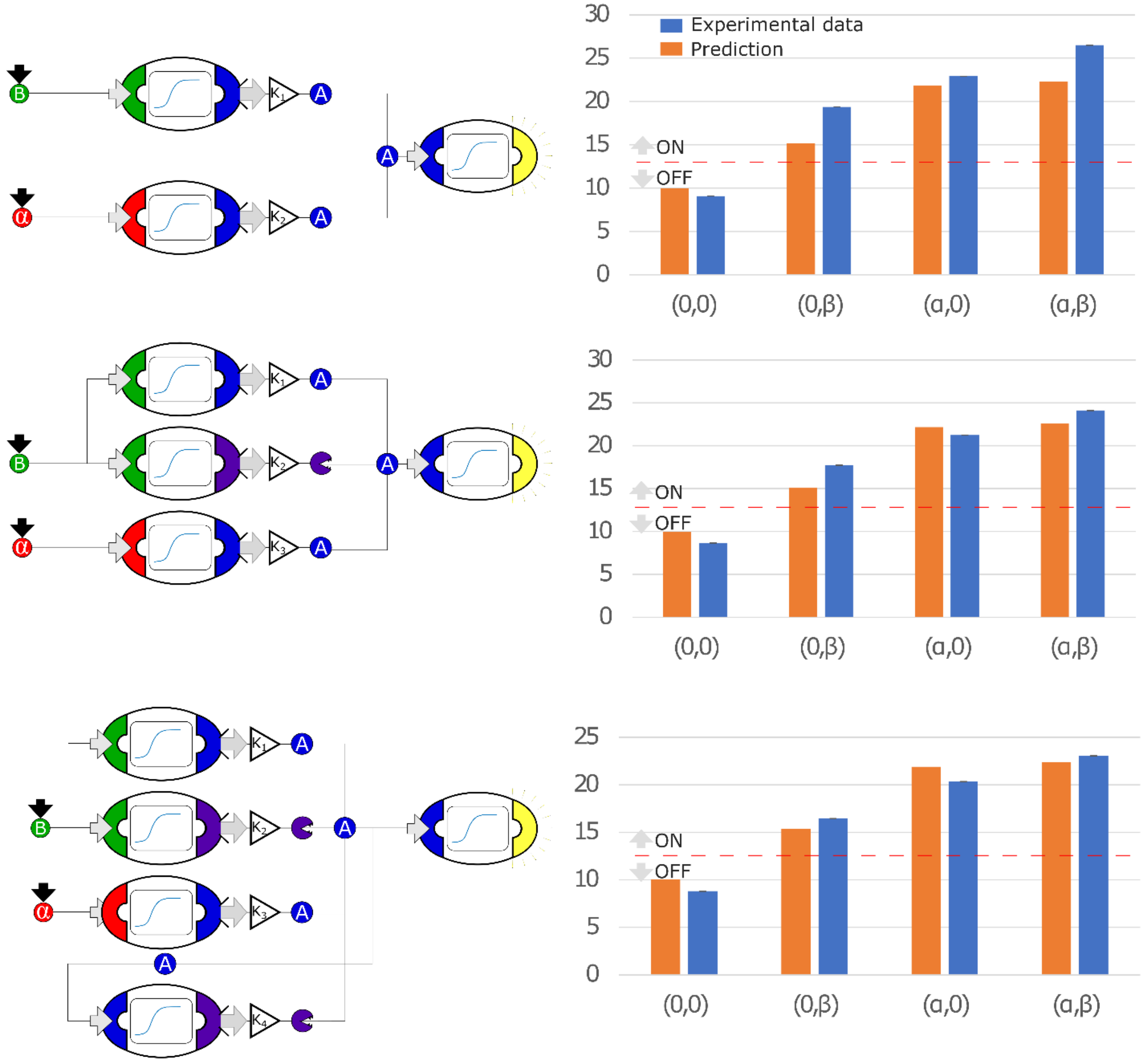

B
α
A
K1
K2
K3
K4
Experimental data
Prediction
ON
OFF
30
25
20
15
10
5
0
(0,0)
(0,β)
(α,0)
(α,β)

### NAND gates

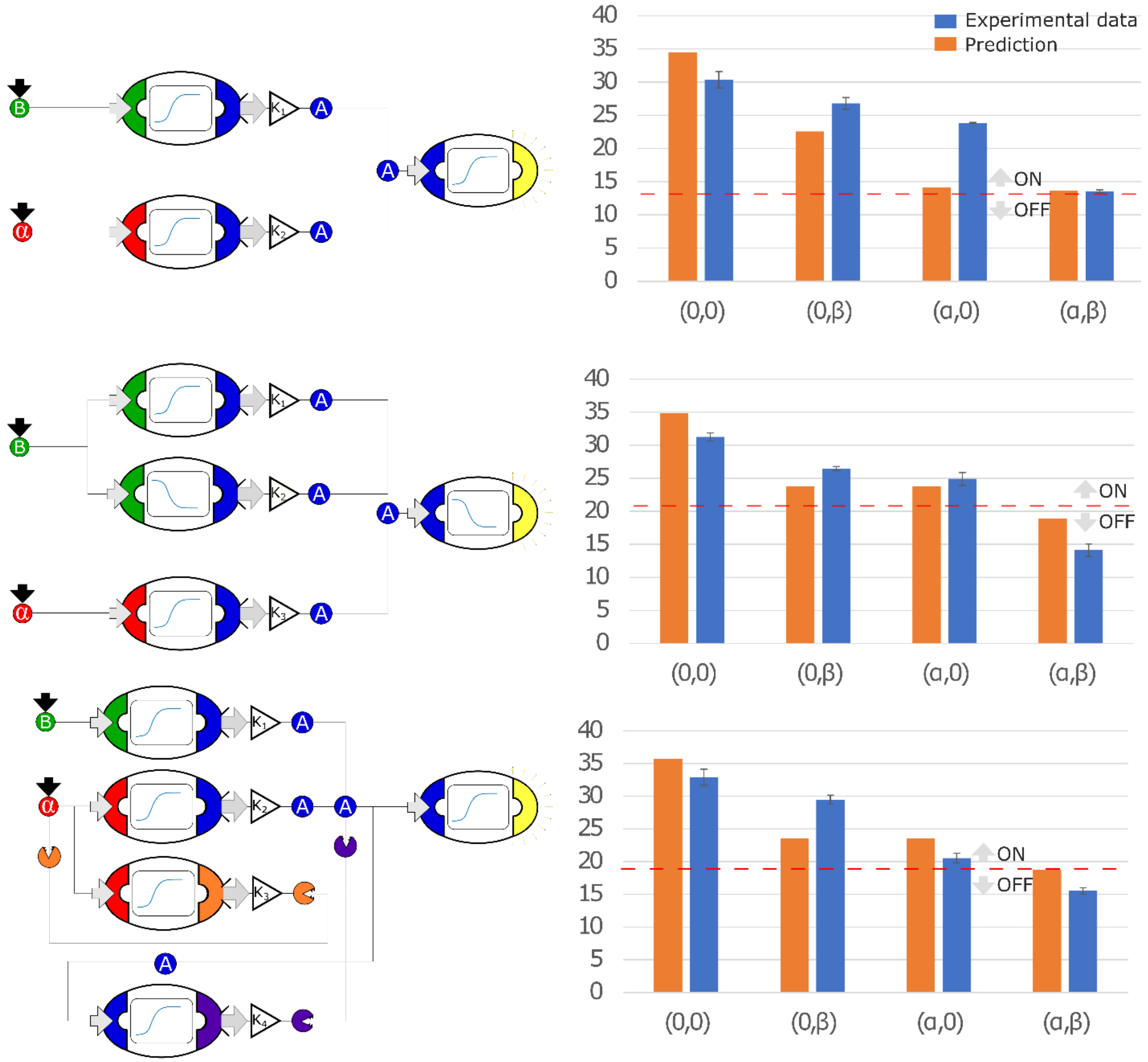

Experimental data
Prediction
ON
OFF
(0,0)
(0,β)
(α,0)
(α,β)
40
35
30
25
20
15
10
5
0
B
α
A
K1
K2
K3
K4

### NOR gates

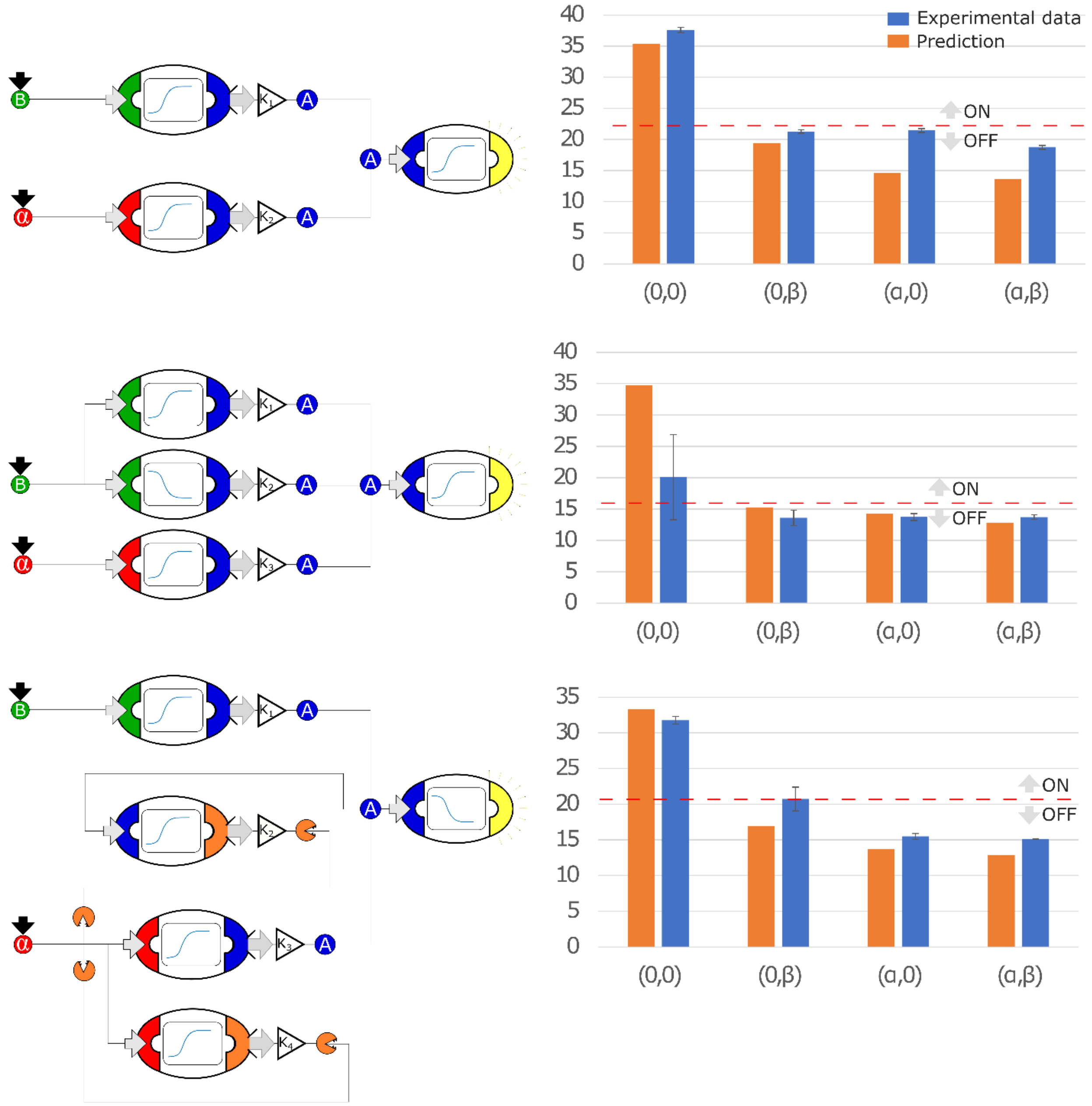


*Figure SI 12:* ***Optimal circuits, simulation and experimental realization of the AND, OR, NAND, NOR gates for 2, 3 and 4-node topologies***

**Table SI 1: all model parameters**

Parameter fit values Table describing parameter estimated for Models 1 in Figure 1. Parameters were estimated using Matlab® *fminsearch* function to minimize the L2 norm of the difference between observation and simulation. Each parameter was estimated three times on three different experimental repeats. Parameter mean and standard deviation are reported in the table against strain # as identified in the strains listed in this study (SI table 1). The parameters used for all the simulation in this study were estimated by fitting the average measurement directly, and it is presented in brackets below the mean±std values.

| | | **Parameter** | | | | | | | | |
|---|---|---|---|---|---|---|---|---|---|---|
| | | $K_1$ | $\delta_1$ | $K_2$ | $\psi$ | n | $\delta_2$ | $K_3$ | $\delta_3$ | b |
| **Strain #** | 1 | 1.32e+06±5.79e+05 (1.76e+06) | 3.41e+06±4.25e+05 (3.13e+02) | 7.08e+05±2.75e+05 (8.29e+05) | 60.52±84.57 (256.87) | 0.59±0.24 (0.89) | 2.46e+05±2.51e+04 (4.41e+5) | 2.50e05±3.52e+04 (2.12e+05) | 5.39±1.81e+94 (7.46e+04) | 864.63±166.77 (867.98) |
| | 2 | 0.65±0.27 (0.69) | 326.27±101.52 (416.02) | 201.66±59.58 (277.36) | 2.07±0.51 (1.79) | 1.01±0.30 (1.011) | 10.99±1.63 (11.40) | 0.0019±7.45e-04 (0.0011) | 0.498±0.019 (0.49) | 0.0044±0.0023 (0.0049) |
| | 3 | 9.64e+04±2.39e+04 (8.95e+04) | 0.092±0.086 (0.082) | 2.53e+07±8.37e+06 (1.73e+07) | 1.097e+07±1.835e+06 (1.24e+07) | 0.83±0.0059 (0.836) | 1.66e+07±2.50e+06 (1.88e+07) | 2.66e+04±4.95e+03 (3.15e+04) | 2.25e+06±4.47e+05 (1.66e+06) | 1.98e+04±5.49e+03 (1.46e+04) |
| | 4 | 5.73±2.35 (6.05) | 51.20±18.34 (59.54) | 4.46±4.15 (10.61) | 0.72±0.17 (0.67) | 0.80±0.0015 (0.80) | 0.79±0.23 (0.82) | 0.0026±0.0028 (3.66e-04) | 1.89±1.59 (1.10) | 0.0054±0.0045 (0.0032) |
| | 5 | 7.49e+05±1.23e+06 (1.06e+03) | 3.19e+05±6.36e+05 (0.245) | 3.06e+06±3.6e+06 (3.47e+05) | 8.54e+05±4.96e+05 (7.08e+05) | 1.60±0.88 (1.04) | 1.77e+06±2.58e+06 (1.56e+05) | 8.036e+04±1.23e+05 (1.62e+04) | 8.20e+06±1.19e+07 (2.15e+06) | 2.36e+04±3.46e+04 (5.96e+03) |
| | 6 | 48.74±10.42 (50.66) | 35.04±10.09 (25.86) | 8.77±2.41 (11.00) | 38.14±12.70 (57.12) | 1.26±0.031 (1.26) | 87.98±16.44 (110.43) | 0.093±0.02 (0.089) | 0.16±0.0035 (0.16) | 2.13e-04±3.68e-05 (2.155e-04) |
| | 7 | 1.18±0.6 (0.89) | 0.18±0.13 (0.17) | 243.78±62.08 (174.32) | 9.16±1.93 (6.65) | 2.12±0.26 (2.18) | 0.58±0.097 (0.56) | 1.82e-04±7.53e-05 (1.5e-04) | 0.63±0.073 (0.55) | 6.32e-04±1.97e-04 (3.77e-04) |

| | | | | | | | | | |
|---|---|---|---|---|---|---|---|---|---|
| **Units** | Molecule NucVol$^{-1}$nM$^{-1}$hour$^{-1}$ | hour$^{-1}$ | Molecule NucVol$^{-1}$ hour$^{-1}$ | Molecule NucVol$^{-1}$ | dimensionless | NucVol Molecule$^{-1}$ hour$^{-1}$ | Fluorescence Arbitrary Units (FAU) hour$^{-1}$ | NucVol Molecule$^{-1}$ hour$^{-1}$ | FAU Molecule NucVol$^{-1}$ hour$^{-1}$ |
| **Description** | Ligand binding affinity of membrane protein(α)/co-factor (IAA)/ TF (β) | Dissociation/degradation/dilution rate of ligand-binder complex | Maximum/minimum transcription rate of TF activator/repressor | Ligand-binder complex concentration producing half occupation | Hill-coefficient | Dissociation rate of DNA-TF | Fluorescence protein accumulation over time normalized to cell size | Fluorescence protein degradation/dilution rate over time | Baseline fluorescence protein accumulation over time normalized to cell size |

Table presenting the parameters estimated for Model 1 for Single-Input/Single-Output strains in Figure 2 and 3. In the first column, the strain # is reported, based on the List of strains used in this study (SI Table 1); the second column reports the sensor strain architecture that was used as baseline but with different output pathway; the parameters here reported refers uniquely to the output pathway: the remaining parameters are identical to the ones estimated for the baseline strain. Each parameter was fitted to data representing the average of three experimental repeats.

| | | | **Parameter** | | |
|---|---|---|---|---|---|
| | | Baseline strain # | $K_3$ | $\delta_3$ | b |
| **Strain #** | 10 | 5 | 566.24 | 0.575 | 55.83 |
| | 11 | 6 | 26.038 | 1.92e-06 | 0.35 |
| | 12 | 6 | 3.48e+03 | 0.16 | 0.21 |
| | 13 | 6 | 121.60 | 0.062 | 0.14 |
| | 14 | 3 | 2.285 | 0.28 | 0.74 |
| | 15 | 6 | 109.96 | 36.71 | 1.60e-04 |
| | 16 | 3 | 1.89 | 1.83e-13 | 0.365 |
| | 17 | 5 | 582.32 | 368.67 | 1.17e-14 |
| | 18 | 4 | 4.09 | 8.74e-11 | 47.78 |
| | 19 | 7 | 2.024e+11 | 4.29e+10 | 1.85e+09 |
| | 20 | 7 | 0.077 | 1.77 | 0.18 |
| | 21 | 1 | 471.73 | 0.41 | 1.34 |
| | 22 | 5 | 419.52 | 2.10e+04 | 2.32e+04 |
| | 23 | 3 | 775.05 | 0.84 | 780.90 |
| | 24 | 4 | 0.0014 | 3.74e+07 | 1.096e+08 |
| | 25 | 1 | 225.51 | 42.46 | 3.14e-06 |

| | | | | |
|---|---|---|---|---|
| Units | | nM hour$^{-1}$ | NucVol Molecule$^{-1}$ hour$^{-1}$ | nM Molecule NucVol$^{-1}$ hour$^{-1}$ |
| Description | | Output protein accumulation over time | Output protein degradation/dilution rate | Baseline output protein accumulation over time |

Table reporting parameter values for the two Multi-Input/Single-Output (MISO) strains in Figure 1E. The parameters were estimated by minimizing the L2 norm of the difference between the simulation of Model 2 (Figure 1) and the average of three experimental repeats. The two MISO strains are labelled 8 or 9 following the table 'List of strains used in this study' (SI Table 1)

| | | Strain # | | | |
|---|---|---|---|---|---|
| | | 8 | 9 | Units | Description |
| Parameter | $K_1$ | 0.0036 | 3.14E-06 | Molecule NucVol$^{-1}$nM$^{-1}$ hour$^{-1}$ | Ligand binding affinity of membrane protein(α)/co-factor (IAA)/ TF (β) |
| | $\delta_1$ | 0.0021 | 6.74E-04 | hour$^{-1}$ | Dissociation/degradation/dilution rate of ligand-binder complex |
| | $K_2$ | 6.92E-05 | 1.34E-19 | Molecule NucVol$^{-1}$nM$^{-1}$hour$^{-1}$ | Ligand binding affinity of membrane protein(α)/co-factor (IAA)/ TF (β) |
| | $\delta_2$ | 0.0047 | 7.46E-05 | hour$^{-1}$ | Dissociation/degradation/dilution rate of ligand-binder complex |
| | $K_3$ | 0.059 | 298.86 | Molecule NucVol$^{-1}$nM$^{-1}$hour$^{-1}$ | Maximum/minimum transcription rate of TF activator/repressor |
| | $\psi_1$ | 0.3 | 1.3 | Molecule NucVol$^{-1}$ | Ligand-binder complex concentration producing half occupation |
| | $n_1$ | 0.31 | 0.003 | dimensionless | Hill coefficient |
| | $\delta_3$ | 3.235 | 574.73 | NucVol Molecule$^{-1}$ hour$^{-1}$ | Dissociation rate of DNA-TF |
| | $K_4$ | 74 | 0.012 | Molecule NucVol$^{-1}$nM$^{-1}$hour$^{-1}$ | Maximum/minimum transcription rate of TF activator/repressor |
| | $\psi_2$ | 1.93 | 0.13 | Molecule NucVol$^{-1}$ | Ligand-binder complex concentration producing half occupation |
| | $n_2$ | 0.85 | 0.6 | dimensionless | Hill coefficient |
| | $\delta_4$ | 58.34 | 0.13 | NucVol Molecule$^{-1}$ hour$^{-1}$ | Dissociation rate of DNA-TF |

| | | | | | |
|---|---|---|---|---|---|
| | $K_5$ | 0.015 | 0.015 | FAU hour$^{-1}$ NucVol Molecule$^{-1}$ | Background fluorescence protein accumulation over time normalized to cell size |
| | $\phi_1$ | 0.32 | 0.043 | FAU hour$^{-1}$ | Fluorescence protein synthesis by the activator over time normalized to cell size |
| | $\phi_2$ | 0.87 | 2.09E+03 | dimensionless | Repressor molecule number needed to degrade one activating molecule |
| | $\delta_5$ | 0.9 | 0.52 | hour$^{-1}$ | Fluorescence protein degradation/dilution rate over time |
